## supplemental for "THAP9 transposase cleaves DNA via conserved acidic residues in an RNaseH-like domain"

### Supplementary Information

**This PDF file includes:** Figures S1 to S7, Table S1

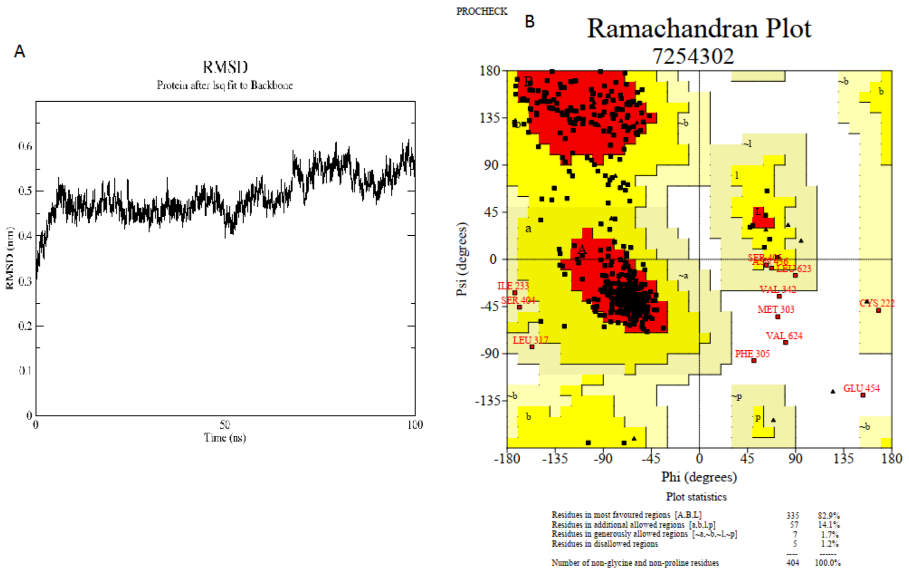


**Fig S1:** RMSD plot and Ramachandran plot for hTHAP9 model after MD simulations.

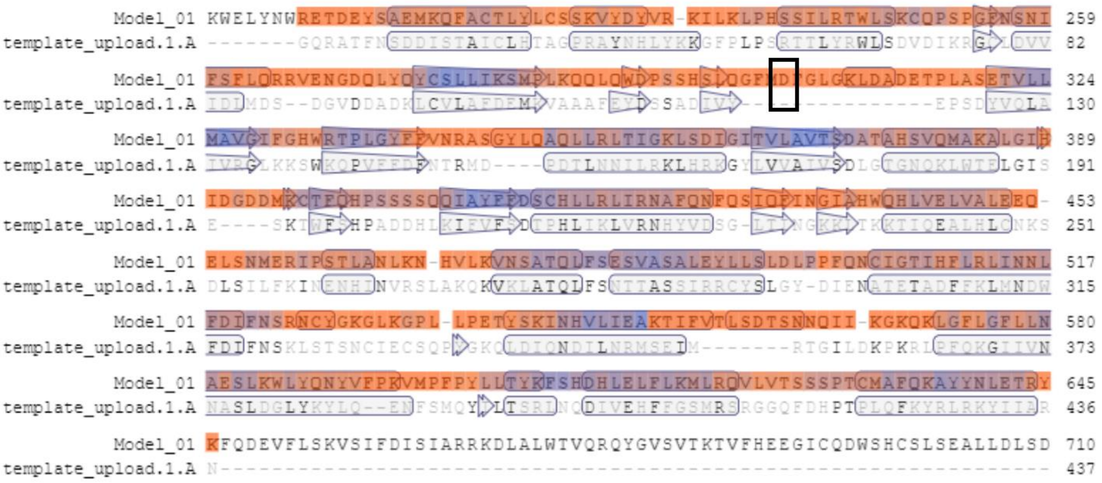


**Fig S2:** SWISS MODEL secondary structure alignment of hTHAP9 (Model_01) homology structure built with DmTNP template (template_upload.1.A). The residues enclosed in horizontal bars represent helical secondary structure and residues enclosed in arrows represent beta sheets. The colour of the sequence represents QMEAN, orange to blue indicates low to high quality estimate (23). D304 of hTHAP9, highlighted in black box, does not align to any structure and ends up in a loop in the final structure. Residue numbers correspond to the amino acid position in the primary sequence of the protein

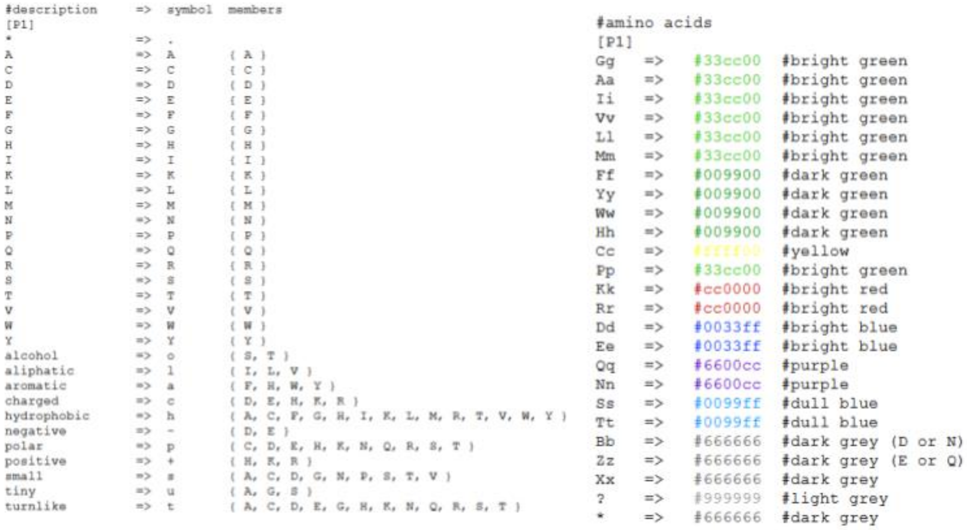

```
#description     =>  symbol  members
[P1]
*                =>  .
A                =>  A       { A }
C                =>  C       { C }
D                =>  D       { D }
E                =>  E       { E }
F                =>  F       { F }
G                =>  G       { G }
H                =>  H       { H }
I                =>  I       { I }
K                =>  K       { K }
L                =>  L       { L }
M                =>  M       { M }
N                =>  N       { N }
P                =>  P       { P }
Q                =>  Q       { Q }
R                =>  R       { R }
S                =>  S       { S }
T                =>  T       { T }
V                =>  V       { V }
W                =>  W       { W }
Y                =>  Y       { Y }
alcohol          =>  o       { S, T }
aliphatic        =>  l       { I, L, V }
aromatic         =>  a       { F, H, W, Y }
charged          =>  c       { D, E, H, K, R }
hydrophobic      =>  h       { A, C, F, G, H, I, K, L, M, R, T, V, W, Y }
negative         =>  -       { D, E }
polar            =>  p       { C, D, E, H, K, N, Q, R, S, T }
positive         =>  +       { H, K, R }
small            =>  s       { A, C, D, G, N, P, S, T, V }
tiny             =>  u       { A, G, S }
turnlike         =>  t       { A, C, D, E, G, H, K, N, Q, R, S, T }
```

```
#amino acids
[P1]
Gg    =>    #33cc00    #bright green
Aa    =>    #33cc00    #bright green
Ii    =>    #33cc00    #bright green
Vv    =>    #33cc00    #bright green
Ll    =>    #33cc00    #bright green
Mm    =>    #33cc00    #bright green
Ff    =>    #009900    #dark green
Yy    =>    #009900    #dark green
Ww    =>    #009900    #dark green
Hh    =>    #009900    #dark green
Cc    =>    #ffff00    #yellow
Pp    =>    #33cc00    #bright green
Kk    =>    #cc0000    #bright red
Rr    =>    #cc0000    #bright red
Dd    =>    #0033ff    #bright blue
Ee    =>    #0033ff    #bright blue
Qq    =>    #6600cc    #purple
Nn    =>    #6600cc    #purple
Ss    =>    #0099ff    #dull blue
Tt    =>    #0099ff    #dull blue
Bb    =>    #666666    #dark grey (D or N)
Zz    =>    #666666    #dark grey (E or Q)
Xx    =>    #666666    #dark grey
?     =>    #999999    #light grey
*     =>    #666666    #dark grey
```

**Fig S3**: P1 colour palette for multiple sequence alignment viewed in MView.

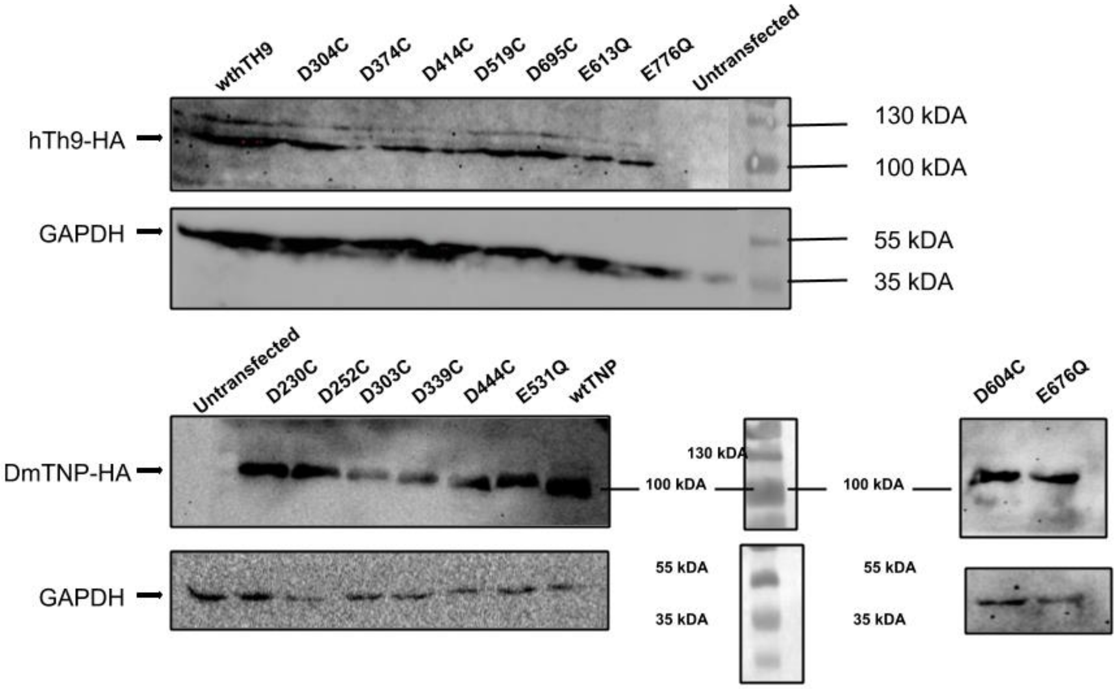


**Fig S4:** Expression levels of mutant and wild-type hTHAP9 and DmTNP (~100 kDa, C-terminal HA tagged) in the DNA excision assay as shown by western blot analysis (anti-HA). GAPDH (mol weight ~36 kDa) was used as a loading control. Negative control (HEK293 cells alone)

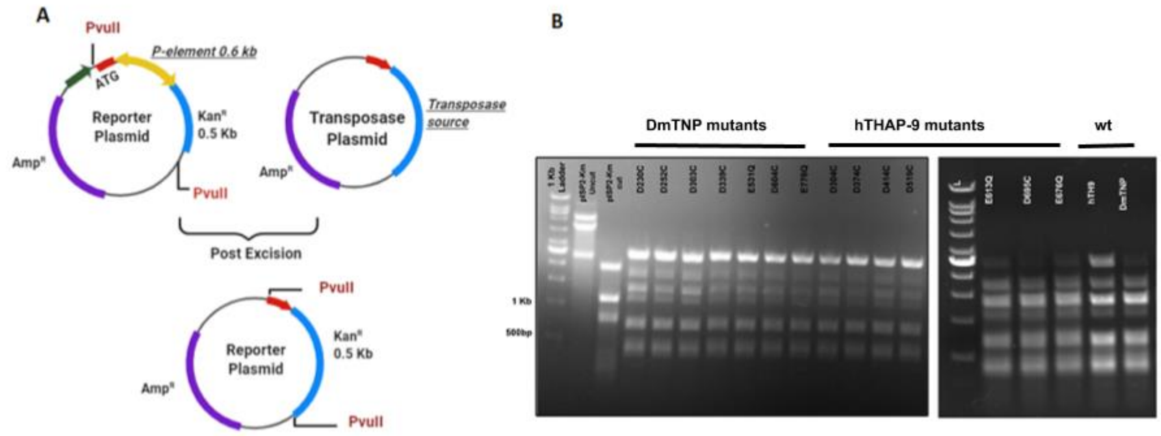


**Fig S5** (A) Schematic representation of PvuII digestion assay of recovered reporter plasmid (obtained after transformation of bacteria with plasmid DNA isolated from HEK 293 cells transfected with DmTNP, hTHAP9 or the acidic residue mutants) (B) Agarose gel image of reporter plasmid after restriction digestion with PvuII enzyme

A. **P element insertion site in pISP2/Km**

2232 **8bp TSD**

ATGACCATGATTACGAATTCCCGGGGATCTGCCGCTGGACTACGTGG **GTCTGGCC**

**17nt P element 5' end** **0.6 kb** **17nt P element 3' end** 2901 **8bp TSD**

CATGATGAAATAACATA- Pelement- TATGTTATTTCATCATG **GTCTGCC**ATTCTCATC

(2273-2289)

B. **Donor site repair**

ATGACCATGATTACG----------------------------------------------------CACTGGCCGTCGTT

2232 4088

Δ 40 upstream Downstream rearrangement

**Fig S6. DNA sequence analysis of P element excision products** (A) Schematic of P element insertion site within pISP2/Km plasmid. The 8bp target site duplications (TSDs) flanking the P element site insertion (red) are underlined and bold. The underlined ATG, upstream of P element insertion is the start codon of LacZ gene in pISP2-Km (20). (B) Typical pattern of donor site repair junction obtained by sequencing pISP2-Km plasmids recovered from individual kanamycin-resistant bacterial colonies (both wildtype and mutant DmTNP and hTHAP9). The red dashed lines represent the excised P element. Blue numbers represent the nucleotide position within pISP2/Km reporter plasmid. It is observed that the P element excision and subsequent repair often involve DNA rearrangements and extensive deletions, as observed before (16).

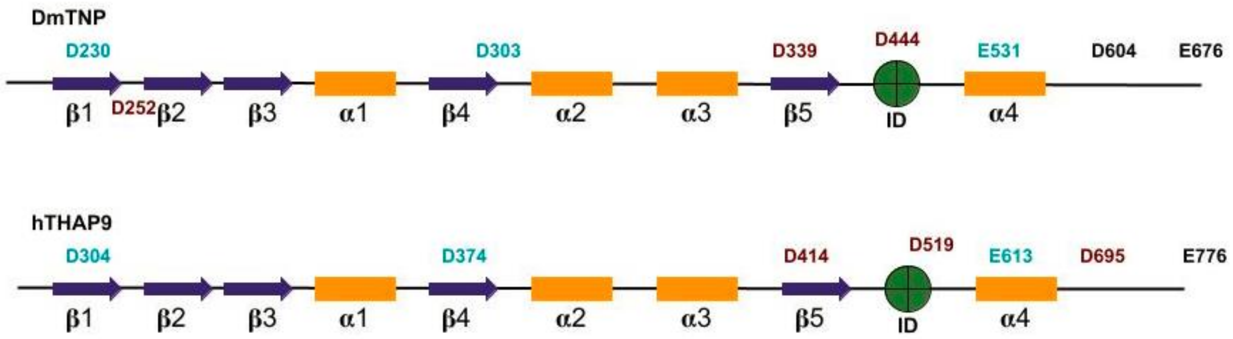


**Fig S7: Summary of catalytic mutants in Drosophila and human THAP9**. DDE motifs for each homolog are represented in bold (cyan) above its putative location. Other conserved residues mutated for the study are marked in black and brown color. The mutations that result in lower excision activity of the transposases are represented in brown. Blue arrows indicate beta sheets and yellow bars represent alpha helices. Insertion domains (ID) for each homolog [DmTNP(residues 339-528) (12), hTHAP9 (residues 415-604), Pdre2 (residues 475-670) are shown in green circles.

**Table S1**: Relative DNA excision and DNA integration activities observed in case of all the mutants in plasmid-based assays.

| | **Mutants** | **% Relative excision activity** | **% Relative Integration activity** |
|---|---|---|---|
| DmTNP | D230C | 12.7 | 19 |
| | D252C | 7.83 | 31 |
| | D303C | 14.02 | 18.33 |
| | D339C | 23.65 | 76.33 |
| | D444C | 0 | 7.33 |
| | D604C | 12.64 | 63.33 |
| | E531Q | 10.99 | 3 |
| | E676Q | 49.6 | 76.66 |
| | WT DmTNP | 100 | 100 |

| | | | |
|---|---|---|---|
| | pISP2-Km | 0 | - |
| | Cg4-Neo | - | 8 |
| | pCDNA 3.1 | - | 0 |
| | pBluescript | - | 0 |
| **hTHAP9** | D304C | 18.19 | 27 |
| | D374C | 26.07 | 5.5 |
| | D414C | 58.03 | 43.5 |
| | D519C | 10.26 | 14 |
| | E613Q | 10.26 | 7.5 |
| | D695C | 141.47 | 55 |
| | E776Q | 94.75 | 144.5 |
| | WT hTHAP9 | 100 | 100 |
| | pISP2-Km | 0 | - |
| | pBluescript | - | 0 |
| | Cg4-Neo | - | 8 |
| | pCDNA 3.1 | - | 0 |
